# Validation of Reference Genes for RT-qPCR Relative Expression Analysis in Pre-Adult Stages of *Taenia solium*

**DOI:** 10.1101/2022.03.22.485324

**Authors:** David Castaneda-Carpio, Jose Maravi, Renzo Gutierrez-Loli, Valeria Villar, Juan Blume La Torre, Segundo W. Del Aguila, Cristina Guerra-Giraldez

**Affiliations:** Laboratorio de Proliferación y Regeneración Celular. Laboratorios de Investigación y Desarrollo, Facultad de Ciencias e Ingeniería. Universidad Peruana Cayetano Heredia. Lima, Peru

**Author notes:** These authors contributed equally to this work (David Castaneda-Carpio, Jose Maravi).

## Abstract

The larvae-to-adult development of the zoonotic parasitic tapeworm *Taenia solium* involves significant but often clinically overlooked events crucial in cestode biology. The early-adult stages can be studied in vitro, providing a valuable model to examine scolex evagination, strobilation, and worm development. Without a stage-specific transcriptome, postgenomic data exploration followed by single-gene relative expression analysis using RT-qPCR (reverse transcription-quantitative PCR) are effective strategies to study gene regulation during parasite development. However, achieving accurate comparisons with this approach requires the validation of an endogenous reference gene (RG).

To address this, we analyzed the expression stability of 17 candidate reference genes (RGs), representing various biological processes, in the context of the in vitro-induced early adult stages of *T. solium* larvae (cysts). RT-qPCR of the candidate RGs was performed on different stages, defined by distinct morphology in culture, and gene expression stability was comprehensively analyzed using the RefFinder tool. Genes *pgk1, bact1, mapk3, tbp, rpl13*, and *cox1* were ranked as the most stable and were used to normalize the expression of *h2b* and *wnt11a*, which are involved in proliferation and strobilation processes in parasitic tapeworms. This study represents the first attempt to identify reliable normalization standards for transcript analysis in the genus *Taenia*.

## INTRODUCTION

The genus *Taenia* includes over a hundred species of tapeworms, parasites of wildlife and livestock. Three among these, *T. solium, T. saginata*, and *T. asiatica*, cause zoonotic infections, leading to taeniasis and cysticercosis. These helminthiases are part of the neglected tropical diseases prioritized by the WHO^1^.

Cysticercosis can cause life-threatening conditions, particularly in poor, remote areas of developing countries in Africa, Asia, and Latin America, where it impacts the health and livelihoods of rural farming communities. The latest WHO report identified *T. solium* as a leading cause of death from food-borne diseases, resulting in a total loss of 2.8 million disability-adjusted life-years (DALYs)^2,3^.

*T. solium* has a complex life cycle involving an intermediate host (pig) that harbors the larvae in its tissues (cysticercosis), and a definitive host (human) that carries the adult tapeworm in its intestines (taeniasis). The tapeworm carrier spreads thousands of microscopic parasite eggs through feces containing infective hexacanth embryos or oncospheres. Upon ingestion by free-roaming pigs or other humans, active embryos are released from their eggshells and cross the intestinal epithelium, spreading through the bloodstream to skeletal muscle, heart, and nervous system, where they lodge and develop into vesicular larvae (cysts)^4^. Poorly cooked pig meat may thus contain viable cysts. The gastric and bile acids in the human digestive tube activate these, and the already formed scolex inside the vesicle evaginates to attach to the intestinal wall. There, the parasite grows into an adult tapeworm by producing a strobilus of continuous proglottids that will carry new eggs when mature^4,5^.

Due to the complexity of the host-parasite interactions, the complete life cycle of *T. solium* has yet to be reproduced in the laboratory. However, observations in experimental or naturally infected animals have produced valuable knowledge about the oncosphere-to-cyst development and cyst-associated immunopathology in several tissues^6,7^. There is much less information related to cyst-to-adult development, partly because taeniasis has generally mild or non-specific symptoms^4,8^. Nonetheless, studying the stages of taeniasis is necessary to understand endemicity’s critical factors. Notably, a tapeworm carrier is the leading risk factor for acquiring cysticercosis, spreading the infective stage and perpetuating the parasite’s life cycle.

Cyst-to-adult development involves several processes essential in cestode biology, associated with the scolex’s evagination and further strobilation^9^. The early adult stage in *T. solium* can be addressed in vitro, starting from chemically^10^ or enzymatically treated cysts^11^.

With the publication of the *T. solium* genome^12^, the first partial cyst transcriptomes obtained by RNA-seq^13,14^, and the recent transcriptome of the abnormal form of the cyst found in human subarachnoid neurocysticercosis^15^, a significant amount of data is currently available to study stage-specific events in the parasite’s development^16^. Moreover, whole transcriptome analyses in other tapeworms (including families Taeniidae – genera *Taenia* and *Echinococcus*^17-21^, Hymenolepididae^22^, and Mesocestoididae^23^) have led to the identification of conserved pathways among cestodes that complement previous datasets.

RT-qPCR helps analyze single-gene expression and compare experimental conditions due to its high sensitivity and specificity. Validating the expression stability of candidate reference genes (RGs) is thus critical to normalize data when studying any particular species or stages with RT-qPCR^24^. Although a few analyses for expression stability of RGs have been done in other cestodes^25,26^, a similar validation for the pre-adult stages of *T. solium* is lacking. Previous RT-qPCR comparisons used canonical housekeeping gene targets such as *gapdh* (for the enzyme Glyceraldehyde-3-Phosphate Dehydrogenase)^27^.

In this study, a representative set of 17 transcripts involved in key biological processes – such as gene transcription, translation and protein synthesis, protein folding and quality control, signal transduction, cytoskeleton and cell structure, energy metabolism, and mitochondrial respiration – were selected as candidate RGs for the in vitro-induced pre-adult stages of *T. solium*. The expression stability of these genes was analyzed in cultivated cysts, classified into three morphological stages corresponding to different moments in the process of scolex evagination. The RefFinder integrative tool, which combines the statistical algorithms geNorm, NormFinder, BestKeeper, and the deltaCT method, was used for the analysis.

Finally, to validate the reliability of the top-ranked candidates, these were used to normalize the expression of two informative genes: *h2b*, involved in cell proliferation during scolex evagination in tapeworms, and *wnt11a*, a morphogen associated with posterior development in flatworms that acts as an effector in strobilation.

## METHODS

### Selection of candidate RGs and primer design

Primers were designed using Primer3^28^ and BLAST® software based on the 17 mRNA sequences of *T. solium* candidate RGs retrieved from GeneDB databases. The basic parameters used for primer design were as follows: primer length between 18-21 bp, GC content between 45-55%, melting temperature between 55-60 °C, and amplicon length between 60-200 bp. Every primer was 100% complementary to the expected primer binding sites, except for the COX1 primers, which had less (97%) complementarity to the used mRNA sequence. The details of each selected candidate gene and the biological features of each primer set are shown in Table 1.

**Table 1.**
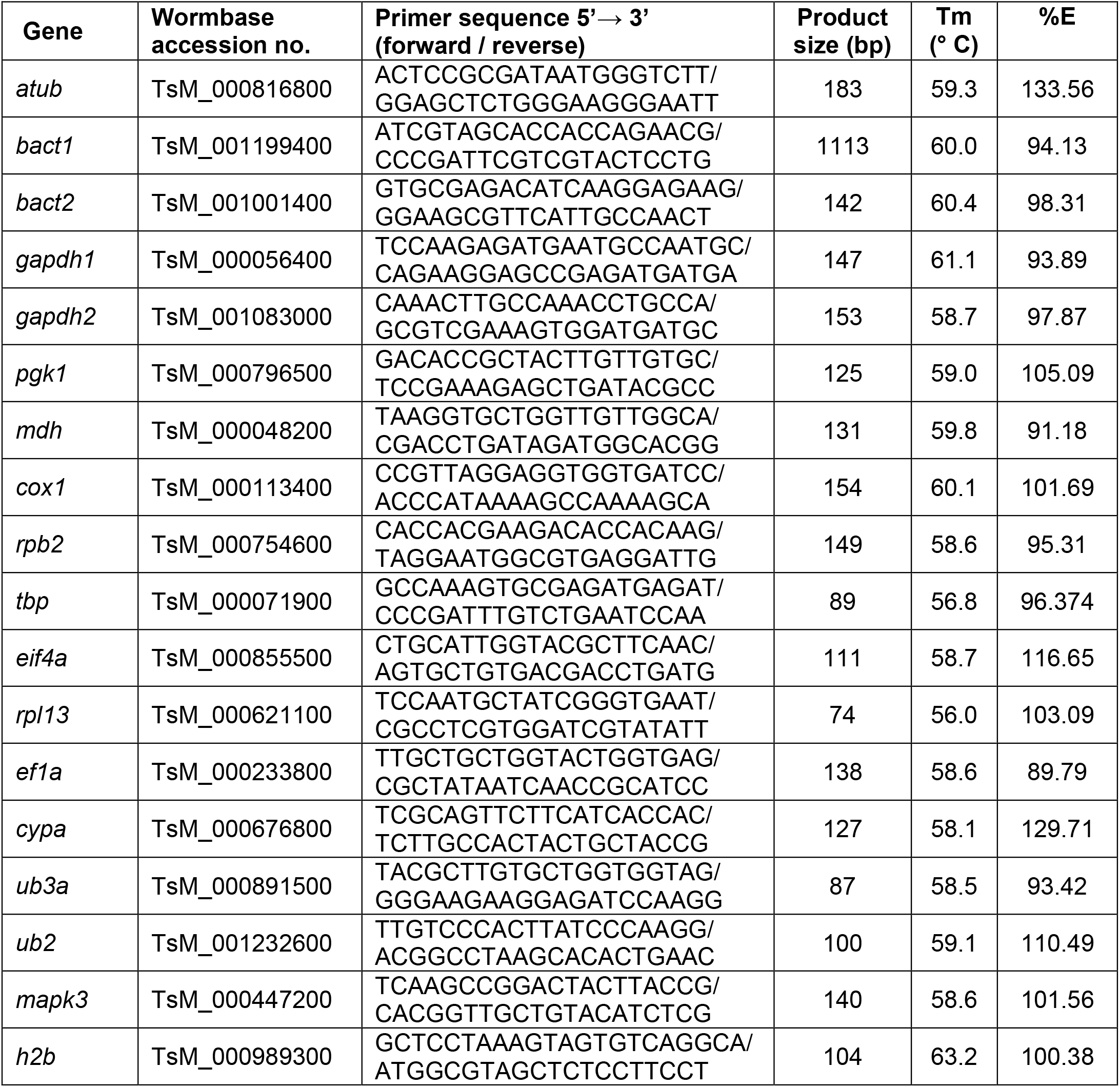
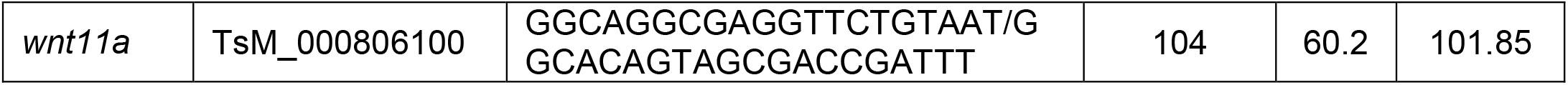
Primer sequences and RT-qPCR amplification features of 17 candidate RG and both target genes.

### Parasite collection and *in vitro* evagination

*Taenia solium* cysts were collected from the muscular tissue of a naturally infected pig, as determined by a positive tongue examination, from a farm at Huancayo, Junin, Peru. Only undamaged and complete cysts were selected and placed in phosphate buffered-saline (PBS; 0.1 M, pH 7.4) supplemented with gentamicin (20 µg/mL) and benzylpenicillin (10 µg/mL) for transport.

The evagination of scolices was induced with 0.1% taurocholic acid (TA) (Sigma, St. Louis, MO) diluted in RPMI 1640 medium, supplemented with 2 g/L NaHCO_3_ (Merck®, Burlington, MA). Parasites were cultured in 12-well plates (Corning, Corning, NY) with 2 mL of the described medium at 37°C in a 5% CO_2_ atmosphere for up to 5 days. Scolex evagination was monitored every 12 hours by direct observation to track intermediate stages until full evagination occurred. Consistent collection criteria were established to categorize parasites into three distinct stages: cysts with invaginated scolex (“non-evaginated cysts,” or PRE), cysts with early scolex exposure (“recently evaginated cysts,” or EV), and cysts showing clear strobila (“fully evaginated cysts,” or POST).

After culture, all parasites were washed with PBS to remove residual media. The percentage of evagination, defined as the proportion of parasites with a visibly evaginated scolex, was recorded for each culture condition and parasite group.

### RNA extraction and DNAse treatment

Evaginated and non-evaginated parasites were embedded independently in 1 mL of TRIzol (Invitrogen, Carlsbad, CA) and admixed with a tissue homogenizer tip (Omni International, Kennesaw, GA) on ice. RNA was isolated following the manufacturer’s protocol, considering an extra washing step with 75% ethanol. After the solubilization, whole RNA was treated with 2 U RNAse-free DNAse I (Invitrogen, Carlsbad, CA) for 30 min at 37° C. The DNAse was inactivated with TRIzol, and the RNA was re-purified. RNA yield was measured by spectrophotometry with NanoDrop® (Thermofisher Scientific, Waltham, MA), and approximately 100 ng of RNA were loaded in a 1.5% agarose gel electrophoresis to test its integrity.

### Two-step RT-qPCR reaction

cDNA was generated from 100 ng of RNA of every parasite by reverse transcription with the High-Capacity Reverse Transcription Kit (Thermofisher Scientific, Waltham, MA) in a final volume of 20 μL. The reaction mix was incubated 15 min at 25° C, 2 h at 37° C and 5 min at 85°C, and stored at 4° C until required.

qPCR reactions were done with the PowerUp® SYBR Green Master Mix (Thermofisher Scientific, Waltham, MA). For every 10-μL reaction, forward and reverse primers were added at a final concentration of 500 nM each, and 1 μL of cDNA was used as the template. The cycling conditions were set in the PikoReal™ Real-Time PCR System (Thermofisher Scientific, Waltham, MA) as follows: 2 min at 50°C, 2 min at 95°C; and 40 cycles of 15 s at 95°C and 1 min at 60°C. Every sample was run in triplicates, and a melting curve step was done at the end of every experiment ranging from 60°C to 95°C with increments of 0.2°C. Additionally, the size and specificity of amplification products were confirmed with a 1.5% agarose gel electrophoresis. RG datasets with Cq values >35 were excluded for having low abundance.

### Amplification efficiency and analytical performance

To determine the PCR amplification efficiency for every candidate RG, 5-point calibration curves of cDNA were prepared using five 2-fold, 5-fold, or 10-fold serial dilutions. The linear regression, slopes, and *y*-axis intercepts for every standard curve were calculated using Graphpad Prism® software version 6.01 (Graphpad®, San Diego, CA). Every dilution was tested in triplicate, and amplification efficiency was determined based on the slope of the log-linear portion of the calibration curve (10^−1/slope^ – 1), following the MIQE guidelines^29^.

### Data analysis for gene expression stability

The gene expression stability of the candidate RGs was compared and ranked with different normalization algorithms integrated in the web-based tool RefFinder (available at: https://www.ciidirsinaloa.com.mx/RefFinder-master/). This program integrates the analysis performed by the extensively used computational algorithms geNorm^30^, Normfinder^31^, BestKeeper^32^, and the comparative DeltaCT method^33^. The selection of the most stable RG was based on the recommended comprehensive ranking calculated by the geometric mean of the stability scores obtained from each method.

### Validation of RG in relative quantification

To validate the reliability of the selected RG, the relative expression of *hb2* and *wnt11a* was normalized by the most and the least stable genes obtained by the normalization algorithms. Indeed, both *h2b* and *wnt11a* genes have been reported previously as differentially expressed upon scolex evagination in parasitic tapeworms^34, 35^. Ten biological samples per condition and three technical replicates per sample were set up for this analysis. Fold-change for *h2b*n and *wnt11a* at experimental conditions was analyzed using the 2^−ΔΔCq^ method^36^. The sequence and other technical details for *h2b* and *wnt11a* primer sets are shown in Table 1.

## RESULTS

### *In vitro* parasite evagination and definition of pre-adult stages

We confirmed that taurocholic acid (TA) induces scolex evagination. This was demonstrated through direct and macroscopic observations and by quantifying the percentage of cysts with evaginated scolices. The results showed that after two days in culture, 0.1% TA in the medium led to twice as many cysts with evaginated scolices as the untreated ones (Fig 1B).

**Fig 1.**
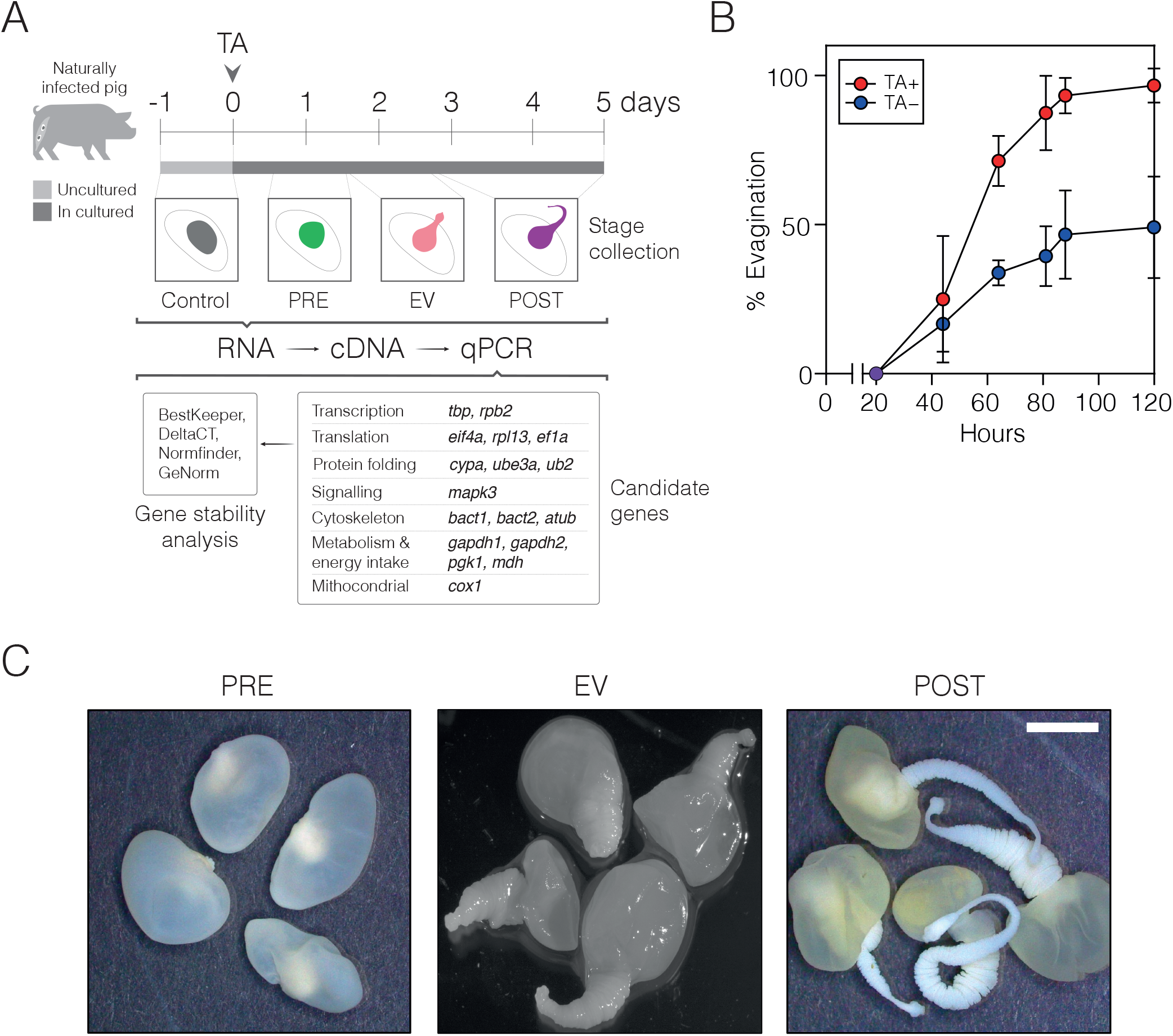
Taurocholic acid (TA) induces in vitro evagination of *T. solium* cysts. (A) Overview of the study design and analysis workflow. (B) Quantification of the percentage of cyst evagination over 120 hours. Each point represents the mean of three independent experiments, with error bars showing the maximum and minimum values. (C) Representative macroscopic images of non-evaginated PRE (left) and POST (right) parasites after 5 days of culture following 0.1% TA induction. Scale bar: 5 mm. These samples were used for subsequent single-gene expression analysis comparisons.

To study the early development of the pre-adult stages, we examined several conditions, including uncultured controls and cysts maintained in vitro with 0.1% TA. Our goal was to identify culture conditions that achieve complete evagination of the scolices, allowing us to establish three *bona fide* categories for comparing single-gene expression against the controls. By five days post-TA induction, the parasites had fully evaginated the scolex, neck, and early strobila from the vesicle (POST). We also included cysts with fully invaginated scolices collected at approximately 4-8 hours in culture (PRE) and cysts collected during the 24-48 hour period (EV), when the parasites had recently evaginated the scolex (Fig 1C).

### RT-qPCR amplification efficiency and product specificity

Primers of 17 candidate RG were used to amplify cDNA templates by qPCR. Agarose gel electrophoresis revealed a single-band amplicon by end-point PCR for each reaction, consistent with the expected product size (Fig 2A). Similarly, all graphs of the negative first derivative of the melting curve showed a single peak, indicating that the primers had strong specificity for the target region and no non-specific amplification occurred (Fig 2D).

**Fig 2.**
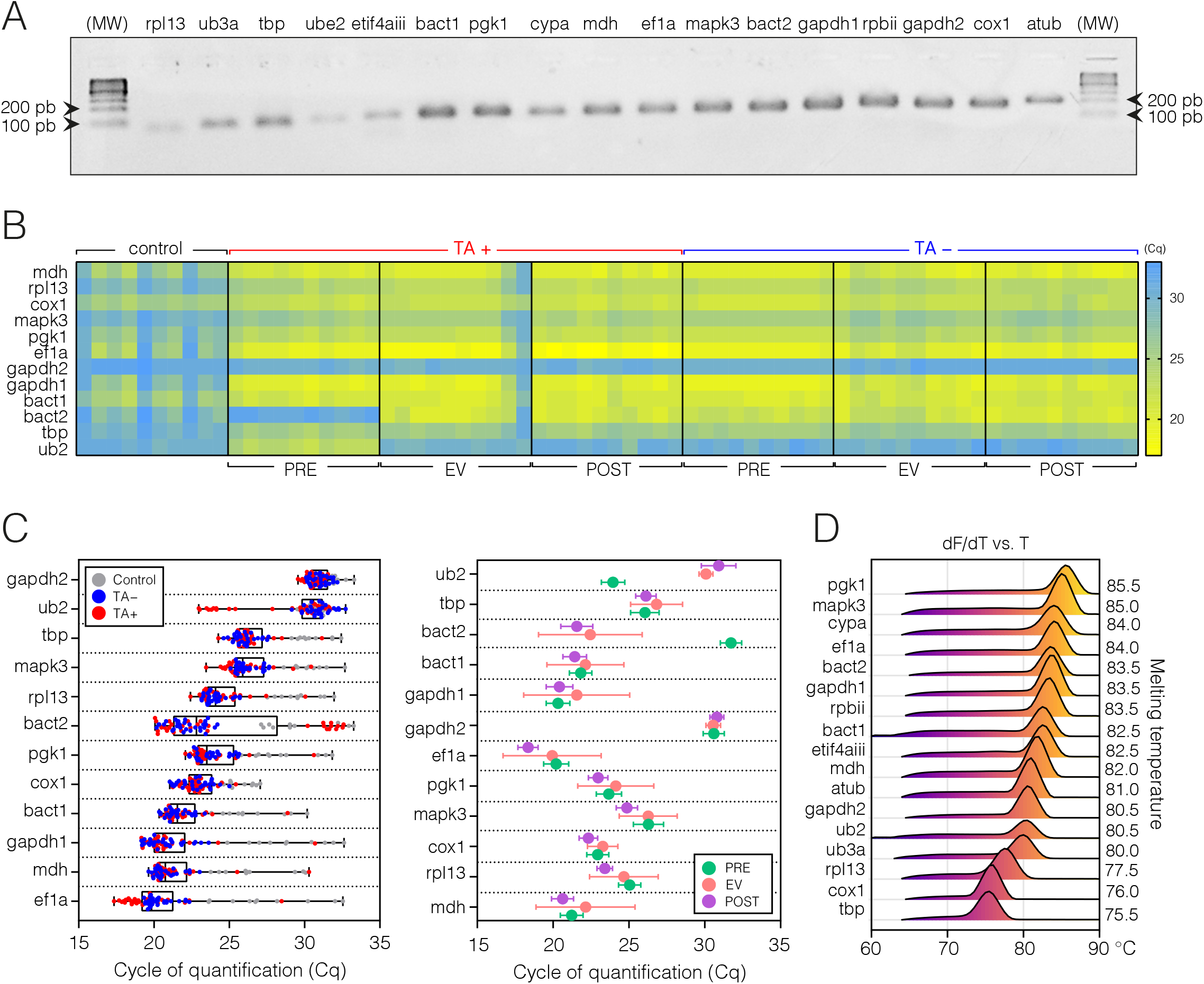
Comprehensive Analysis of qPCR Performance for Candidate Reference Genes. (A) Primer pair specificity for candidate reference genes: End-point PCR products for all candidate RG were observed on gel electrophoresis and arranged according to product size. MW: 100bp DNA Molecular Weight Ladder. (B) Raw Cq values across parasite samples: Comparison of raw Cq values from qPCR amplification across all parasite samples, showing gene expression for each candidate RG. (C) Cq Value distribution based on treatment and morphology: The distribution of Cq values for each candidate RG, grouped by TA treatment (left) and morphological condition of the parasite (right). (D) Ridgeline plot of melting curves for candidate RG: Simplified representation of the melting curves (dF/dT vs. T) obtained from qPCR runs. A single peak was observed for all candidate genes, confirming specific amplification without contamination.

The amplification efficiency for all reactions performed with each primer set reached values above 90%. However, genes *rpbii, etif4aiii, atub, ube3a*, and *cypa* were excluded from downstream analysis because their efficiency values were above 100%, which could lead to inaccurate measurements (Table 1).

### RT-qPCR performance and specificity and RG expression profile

To determine the dispersion of Cq values for the selected candidate RG under conditions representing *T. solium* early-adult stages, initial qPCR runs were performed using cDNA from controls and PRE, EV, and POST parasites. A small Cq value indicated higher gene expression abundance, and vice versa. Mean Cq values for all RG were below 35 cycles, suggesting that transcript abundances were not underrepresented.

A comprehensive heat map of the raw Cq values helped identify parasite samples contributing to variability within their groups. Overall, the samples showed consistent expression levels. Most candidate genes were upregulated due to culture conditions, regardless of AT treatment. Notably, the *ube2* gene, encoding a ubiquitin-conjugating enzyme involved in protein degradation, was specifically upregulated during early culture only with AT treatment (Fig 2B). Similarly, genes *bact2* (beta-actin isoform 2, a cytoskeletal protein) and *ef1a* (Eukaryotic Elongation Factor 1-alpha, crucial for protein synthesis) exhibited distinct AT-dependent differential expression. Interestingly, *gapdh1*, involved in glycolysis and other cellular processes, showed the lowest expression across all conditions and treatments (Fig 2C).

### Comprehensive stability analysis of candidate RG among conditions

RefFinder, a free web tool, combines the results from BestKeeper, geNorm, NormFinder, and the comparative DeltaCT method, using raw Cq values as input to provide a geometric mean ranking of RG stability.

Bestkeeper ranks the candidate RG based on two key indicators: (1) the standard deviation (SD) of the Cq values (genes with SD > 1 are considered unstable), and (2) the Pearson correlation coefficient (r), comparing each gene’s expression to a BestKeeper index derived from the geometric mean of all other genes. Higher correlation (r values close to 1) indicates better stability. geNorm calculates a stability score (M) for each gene, representing the average pairwise variation between that gene and all others being tested. Genes with the highest M values (least stability) are eliminated in a stepwise manner. A lower M value indicates higher stability. A cut-off of 0.15 is used to determine if additional RG are needed. Normfinder ranks RG by calculating their expression stability considering both intra- and inter-group variation. It identifies the genes with the least variation across different conditions or treatments. A stability value below 0.15 is generally considered suitable for normalization. Finally, the deltaCT method compares the relative expression levels of candidate RG pairs across all samples. Gene pairs with the smallest variability (smallest difference in their Cq values) are considered more stable.

The BestKeeper analysis identified *gapdh2, cox1*, and *tbp* as the most stable genes. geNorm found *pgk1* and *bact1* to be equally stable, followed by *rpl13* and *mapk3*. NormFinder ranked *mapk3, tbp*, and *pgk1* as the most stable, while the deltaCT method highlighted *pgk1, rpl13*, and *bact1* as the top stable genes (Fig 3, left). The comprehensive ranking, based on the geometric mean of all analyses, identified *pgk1, bact1, mapk3, tbp, rpl13, cox1*, and *gapdh2* (in descending order) as the most stable genes across all comparisons (Fig 3, right). These genes are essential for key cellular processes, including metabolism, structural integrity, and signal transduction.

**Figure 3.**
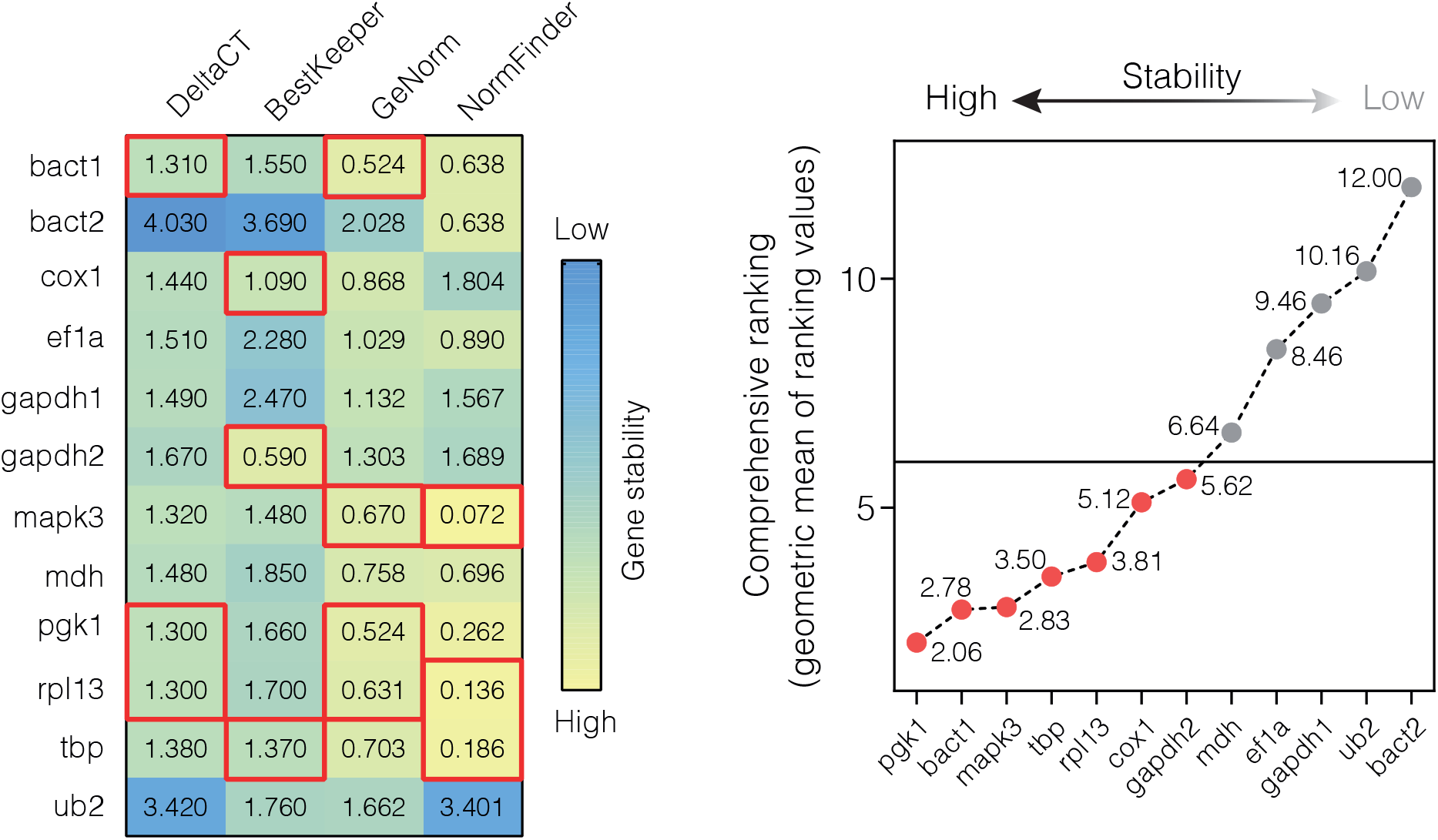
Comprehensive Gene Stability Ranking by RefFinder. The stability scores for candidate RG were analyzed using geNorm, NormFinder, BestKeeper, and the deltaCT method. These results were combined into a comprehensive stability ranking, calculated as the geometric mean of the scores from all four methods. Lower values indicate higher gene stability, reflecting reduced variability.

### Validation of the top-ranked RG

To validate the stability of the selected RG, *h2b* and *wnt11a* were used as target genes under the PRE, EV, and POST early-adult conditions. As previously reported in other parasitic tapeworms, these genes are expected to be upregulated during scolex evagination, as both cell proliferation and strobilation are inherent to this morphogenic process.

After calculating the fold-change values using the 2^-ΔΔCq^ method, the proliferative gene *h2b* showed a 2-to 6-fold increase in expression throughout the early-adult stages compared to the uncultured control. Interestingly, when *h2b* expression was normalized using the *mapk3* and *tbp* RG, the uncultured control groups formed a more consistent cluster with reduced variability. Similarly, *wnt11a* expression was upregulated in cultured parasites, showing a 5-to 10-fold increase when using the different selected RG. The fold-change increase remained within a consistent range for both *h2b* (no values exceeding an 8-fold increase) and *wnt11a* (no values exceeding a 30-fold increase), suggesting that the selected RG exhibit a similar expression pattern, supporting their suitability for normalization (Fig 4A).

**Figure 4.**
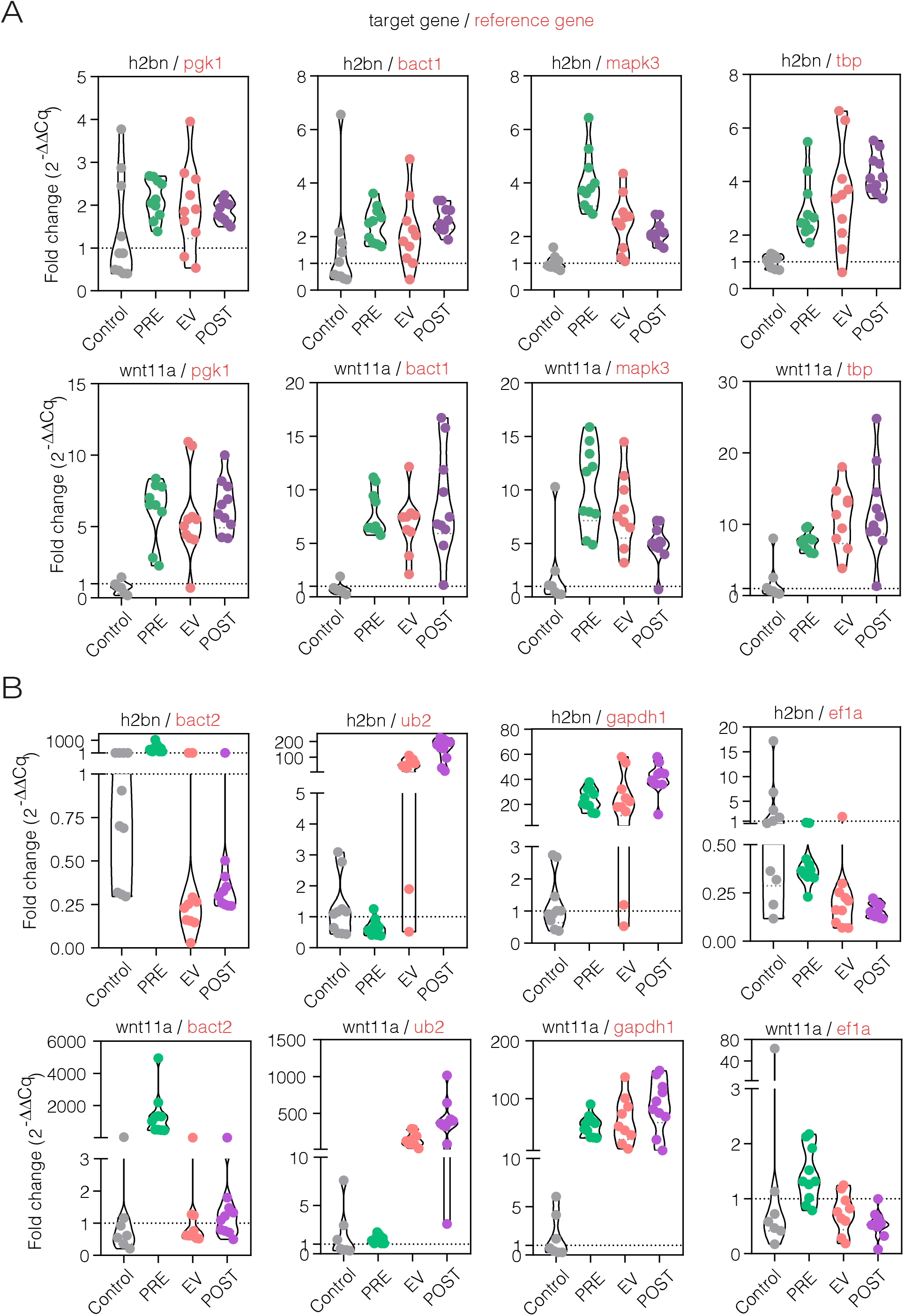
Relative Quantification of *h2b* and *wnt11a* expression using the most and least stable reference genes. The relative expression of *h2b* and *wnt11a*, involved in cell proliferation and strobilation, was calculated using the 2^−ΔΔCq^ method. Gene expression was normalized with (A) the top-ranked and (B) the lowest-ranked reference genes identified through the stability analysis.

In contrast, when the least stable candidates (*bact2, ub2, gapdh1*, and *ef1a*) were used for normalization, the expression levels of *h2b* and *wnt11a* varied significantly across treatments. The *h2b* fold-change fluctuated widely, ranging from 20-to 1000-fold, depending on the gene used as reference. Similarly, *wnt11a* showed highly variable fold-changes, ranging from 10-to 6000-fold, indicating that these unstable genes lead to inconsistent normalization.

Notably, a common feature of the fold-change values obtained for *h2b* and *wnt11a* across the different stages is their inconsistency, displaying a bimodal distribution in gene expression. The values range from below the detection limit (<1, indicating downregulation) to significant increases in expression, as seen when *bact1* or *ef1a* are used for normalization (Fig 4B).

## DISCUSSION

Like in other neglected infectious diseases, the biology of *T. solium* is largely unexplored. This field holds crucial information for better understanding interactions with the hosts, developmental changes through life cycles, and the associated disease endemicity. In the absence of stage-specific transcriptomes, single-gene expression analysis is a powerful approach to exploring the roles of genes of interest under well-defined biological conditions. The controlled in vitro evagination of cysts in the presence ofTA provides a valuable model for studying early-adult development. The normalization of gene expression data often relies on canonical RG that have not been properly validated for this organism or its (yet not described) developmental stages.

In our analysis, we selected a group of genes involved in essential eukaryotic housekeeping processes. Some of them have previously been analyzed for stability in closely related cestodes like *Echinococcus sp*. They are also widely used for normalization in relative quantification studies across various organisms. The comprehensive ranking of gene stability highlighted *pgk1, bact1, mapk3, tbp, rpl13, cox1*, and *gapdh2*.

PGK1 is a glycolytic enzyme. Due to the stability of glycolytic processes across many cell types, it is frequently selected as a housekeeping gene for normalization in qPCR and gene expression analyses in cell lines or animal tissues^37^. However, no similar analyses have been performed in free-living or parasitic worms, suggesting that PGK1 expression may be upregulated (and maintained) in glucose-rich environments, such as those provided by culture media.

While genes involved in signal transduction, such as *mapk3*, are typically considered highly variable, a small study on *H. microstoma* found that cAMP, a key second messenger in signaling pathways, was consistently expressed along the anteroposterior gradient of the adult body^38^. This suggests that the cellular signaling pathways may be stably activated in response to the culture conditions.

Although *gapdh* is commonly used to normalize gene expression, it plays several roles beyond its well-known function in the glycolytic pathway. For instance, it helps maintain cellular iron homeostasis^39^ and acts as a chaperone^40^. Furthermore, its expression levels in human cells are not stable^41^ and show greater variability across different tissues compared to other genes^42,43^. Consequently, previous studies have raised concerns about using GAPDH as a reference gene in qPCR experiments^44^.

Moreover, gapdh has undergone gene duplication events throughout evolution, resulting in different isoforms that may have distinct roles. As a result, the expression of gapdh isoforms can vary depending on the tissue^45^.

Our comprehensive analysis included the isoforms *gapdh1* and *gapdh2*, but neither demonstrated high stability. Although the Bestkeeper algorithm identified *gapdh2* as the most stable candidate, likely due to its low SD across all samples and conditions, other algorithms ranked gapdh2 lower. Additionally, gapdh exhibited the highest raw Cq values among the seventeen initial reference genes, suggesting low expression levels in parasite tissues, making it unsuitable for relative expression analyses. Notably, when gapdh1 was used to normalize the expression of h2b and wnt11a, the resulting fold change values were significantly overestimated compared to those obtained with the top-ranked reference genes.

Some of the top-ranked genes in our analysis, such as *tbp, rpl13*, and *bact1*, were also previously reported as among the most stable in a similar study on *Echinococcus sp*. protoscoleces that were activated in vitro with pepsin^26^. This suggests that, despite being based on a limited number of representative elements, the expression profiles may share common features across species within the phylum Platyhelminthes.

In contrast, one of our top-ranked genes, *tbp*, was identified in a recent study focused on reference gene selection in the free-living echiuran worm *Urechis unicinctus*. Based on transcriptome data, this study suggested that *tbp* is among the least stable genes^46^. However, the genera *Urechis* and *Taenia* occupy distant evolutionary branches within the animal kingdom. Echiurans belong to the Phyllum Annelida, significantly divergent from the Platyhelminths.

Finally, our in vitro design provides a versatile tool for exploring and analyzing the expression of genes of interest throughout the early adult stages during chemically induced evagination of *T. solium*. However, a significant limitation is that our determinations were done on whole-cyst RNA preparations. As seen in other parasitic cestodes, several genes exhibit discrete expression patterns along the adult worm’s body. Techniques such as in situ hybridization have revealed the presence of regional expression gradients, which may be restricted to or extend across anatomical regions like the scolex, the “transition zone” between the neck and the nascent strobila, as well as the immature and mature proglottids^34,35^. Therefore, depending on the specific goals, a more refined set of criteria should be established, such as preparing RNA from carefully selected anatomical regions, to improve the accuracy of the analysis. Nevertheless, this would be difficult on cysts with invaginated scolices (our PRE parasites).

Taken together, the results of this work validated that *pgk1, bact1, mapk3, tbp, rpl13*, and *cox1* could be suitable RG for normalizing single-gene relative expression qRT-PCR data in the context of *T. solium* cyst evagination. This study provides a basis for future studies of gene function in *T. solium* early-adult development.

## ACKNOWLEDGMENTS

We thank Luz Moyano and Ricardo Gamboa (Global Health Tumbes, UPCH) for providing viable *T. solium* cysts; David Duran (Instituto de Medicina Tropical “Alexander von Humboldt”, UPCH) for his kind assistance with gene expression equipment; Vanessa Arevalo (Laboratorio de Ecotoxicología, UPCH) for her assistance with macro photography of cultured parasites, and Francisco Villafuerte, Daniela Bermúdez and Rómulo J. Figueroa (Laboratorio de Fisiología Comparada, UPCH) for sharing their facilities for parasite culture. RGL and JBLT were supported by a training grant from NIH-Fogarty, USA (2D43TW007120-11A1). DC, JM, and VV were supported by a grant from the Peruvian Programa Nacional de Investigación Científica y Estudios Avanzados, Prociencia (PE501079376-2022), which also funded the completion of this work.

## AUTHOR CONTRIBUTIONS

RGL and JBLT conceptualized and designed this work. DC, JM, VV, JBLT, and SDA performed experiments. RGL, DC, and JBLT performed all data analysis. RGL wrote the manuscript with input from all authors, and CGG finished the critical revision of the article.

## REFERENCES

1. World Health Organization. Ending the neglect to attain the Sustainable Development Goals: A road map for neglected tropical diseases 2021–2030. (2020). Available at: https://www.who.int/publications/i/item/9789240010352. (Accessed: 15th February 2021)

2. World Health Organization. WHO estimates of the global burden of foodborne diseases. (2015). Available at: https://www.who.int/foodsafety/publications/foodborne_disease/fergreport/en/. (Accessed: 15th February 2021)

3. World Health Organization. Taeniasis/cysticercosis. (2022). Available at: https://www.who.int/news-room/fact-sheets/detail/taeniasis-cysticercosis. (Accessed: 15th February 2021)

4. Garcia, H. H., Gonzalez, A. E. & Gilman, R. H. Taenia solium cysticercosis and its impact in neurological disease. Clinical Microbiology Reviews 33, (2020).

5. Flisser, A. et al. Taenia solium: Current understanding of laboratory animal models of taeniosis. Parasitology 137, 347–357 (2010).

6. Arora, N. et al. Recent advancements and new perspectives in animal models for Neurocysticercosis immunopathogenesis. Parasite Immunology 39, (2017).

7. Palma, S. et al. In vitro model of postoncosphere development, and in vivo infection abilities of Taenia solium and Taenia saginata. PLoS Negl. Trop. Dis. 13, (2018).

8. Ávila, G. et al. Laboratory animal models for human Taenia solium. in Parasitology International 55, (Parasitol Int, 2006).

9. Koziol, U., Domínguez, M. F., Marín, M., Kun, A. & Castillo, E. Stem cell proliferation during in vitro development of the model cestode Mesocestoides corti from larva to adult worm. Front. Zool. 7, 22 (2010).

10. Rabiela, M. T., Hornelas, Y., Garcia-Allan, C., Rodriguez-del-Rosal, E. & Flisser, A. Evagination of Taenia solium cysticerci: A histologic and electron microscopy study. Arch. Med. Res. 31, 605–607 (2000).

11. Paredes, A. et al. In vitro analysis of albendazole sulfoxide enantiomers shows that (+)-(R)-albendazole sulfoxide is the active enantiomer against Taenia solium. Antimicrob. Agents Chemother. 57, 944–949 (2013).

12. Tsai, I. J. et al. The genomes of four tapeworm species reveal adaptations to parasitism. Nature 496, 57–63 (2013).

13. Almeida, C. R. et al. Transcriptome analysis of Taenia solium cysticerci using open reading frame ESTs (ORESTES). Parasites and Vectors 2, (2009).

14. Lundström, J., Salazar-Anton, F., Sherwood, E., Andersson, B. & Lindh, J. Analyses of an expressed sequence tag library from Taenia solium, cysticerca. PLoS Negl. Trop. Dis. 4, 1–8 (2010).

15. Orrego, M. A. et al. Transcriptomic analysis of subarachnoid cysts of Taenia solium reveals mechanisms for uncontrolled proliferation and adaptations to the microenvironment. Sci. Rep. 14(1):11833 (2024).

16. Paludo, G. P. et al. Cestode strobilation: Prediction of developmental genes and pathways. BMC Genomics 21, (2020).

17. Yang, D. et al. Annotation of the transcriptome from taenia pisiformis and its comparative analysis with three taeniidae species. PLoS One 7, (2012).

18. Parkinson, J. et al. Correction: A Transcriptomic Analysis of Echinococcus granulosus Larval Stages: Implications for Parasite Biology and Host Adaptation. PLoS Negl. Trop. Dis. 6, (2012).

19. Wu, X. et al. Detailed Transcriptome Description of the Neglected Cestode Taenia multiceps. PLoS One 7, (2012).

20. Zhang, S. Comparative transcriptomic analysis of the larval and adult stages of Taenia pisiformis. Genes (Basel). 10, (2019).

21. García-Montoya, G. M. et al. Transcriptome profiling of the cysticercus stage of the laboratory model Taenia crassiceps, strain ORF. Acta Trop. 154, 50–62 (2016).

22. Olson, P. D. et al. Genome-wide transcriptome profiling and spatial expression analyses identify signals and switches of development in tapeworms. Evodevo 9, (2018).

23. Basika, T. et al. Transcriptomic profile of two developmental stages of the cestode parasite Mesocestoides corti. Mol. Biochem. Parasitol. 229, 35–46 (2019).

24. Huggett, J., Dheda, K., Bustin, S. & Zumla, A. Real-time RT-PCR normalisation; strategies and considerations. Genes and Immunity 6, 279–284 (2005).

25. Pouchkina-Stantcheva, N. N., Cunningham, L. J. & Olson, P. D. Spatial and temporal consistency of putative reference genes for real-time PCR in a model tapeworm. Mol. Biochem. Parasitol. 180, 120–122 (2011).

26. Espínola, S. M., Ferreira, H. B. & Zaha, A. Validation of suitable reference genes for expression normalization in Echinococcus spp. larval stages. PLoS One 9, (2014).

27. Hou, J. et al. Sequence analysis and molecular characterization of Wnt4 gene in metacestodes of Taenia solium. Korean J. Parasitol. 52, 163–168 (2014).

28. Untergasser, A. et al. Primer3-new capabilities and interfaces. Nucleic Acids Res. 40, (2012).

29. Bustin, S. A. et al. The MIQE guidelines: minimum information for publication of quantitative real-time PCR experiments. Clin. Chem. 55, 611–22 (2009).

30. Vandesompele, J. et al. Accurate normalization of real-time quantitative RT-PCR data by geometric averaging of multiple internal control genes. Genome Biol. 3, RESEARCH0034 (2002).

31. Andersen, C. L., Jensen, J. L. & Ørntoft, T. F. Normalization of real-time quantitative reverse transcription-PCR data: A model-based variance estimation approach to identify genes suited for normalization, applied to bladder and colon cancer data sets. Cancer Res. 64, 5245–5250 (2004).

32. Pfaffl, M. W., Tichopad, A., Prgomet, C. & Neuvians, T. P. Determination of stable housekeeping genes, differentially regulated target genes and sample integrity: BestKeeper - Excel-based tool using pair-wise correlations. Biotechnol Lett. 26, 509–515 (2004).

33. Silver, N., Best, S., Jiang, J. & Thein, S. L. Selection of housekeeping genes for gene expression studies in human reticulocytes using real-time PCR. BMC Mol Biol. 7, 33 (2006).

34. Jarero, F. et al. Muscular remodeling and anteroposterior patterning during tapeworm segmentation. Dev Dyn. 1–26 (2024)

35. Rozario, T., Quinn, E.B., Wang, J., Davis, R.E. & Newmark, P.A. Region-specific regulation of stem cell-driven regeneration in tapeworms. Elife. 8:e48958 (2019).

36. Livak, K. J. & Schmittgen, T. D. Analysis of relative gene expression data using real-time quantitative PCR and the 2-ΔΔCT method. Methods. 25, 402–408 (2001).

37. Schwarz, A. P. et al. Reference gene expression stability within the rat brain under mild intermittent ketosis induced by supplementation with medium-chain triglycerides. PloS one, 18(2), e0273224 (2023).

38. Pouchkina-Stantcheva, N. N., Cunningham, L. J., & Olson, P. D. Spatial and temporal consistency of putative reference genes for real-time PCR in a model tapeworm. Mol Biochem Parasitol. 180(2), 120–122 (2011).

39. Boradia, V. M., Raje, M., & Raje, C. I Protein moonlighting in iron metabolism: glyceraldehyde-3-phosphate dehydrogenase (GAPDH). Biochem Soc Trans. 42(6), 1796–1801 (2014).

40. Sweeny, E. A. et al. Glyceraldehyde-3-phosphate dehydrogenase is a chaperone that allocates labile heme in cells. J Biol Chem. 293(37), 14557–14568 (2018).

41. Zhu, G., Chang, Y., Zuo, J., Dong, X., Zhang, M., Hu, G., & Fang, F. Fudenine, a C-terminal truncated rat homologue of mouse prominin, is blood glucose-regulated and can up-regulate the expression of GAPDH. Biochem Biophys Res Commun. 281(4), 951–956 (2001).

42. Barber, R. D., Harmer, D. W., Coleman, R. A., & Clark, B. J. GAPDH as a housekeeping gene: analysis of GAPDH mRNA expression in a panel of 72 human tissues. Physiol Genomics. 21(3), 389–395 (2005).

43. Radonić, A., Thulke, S., Mackay, I. M., Landt, O., Siegert, W., & Nitsche, A. Guideline to reference gene selection for quantitative real-time PCR. Biochem Biophys Res Commun. 313(4), 856–862 (2004).

44. Ke, L. D., Chen, Z., & Yung, W. K. A reliability test of standard-based quantitative PCR: exogenous vs endogenous standards. Mol Cell Probes. 14(2), 127–135 (2000).

45. Manchado, M., Infante, C., Asensio, E., & Cañavate, J. P. Differential gene expression and dependence on thyroid hormones of two glyceraldehyde-3-phosphate dehydrogenases in the flatfish Senegalese sole (Solea senegalensis Kaup). Gene. 400(1-2), 1–8 (2007).

46. Chen, J., Wang, Y., Yang, Z., Liu, D., Jin, Y., Li, X., Deng, Y., Wang, B., Zhang, Z., & Ma, Y. Identification and validation of the reference genes in the echiuran worm Urechis unicinctus based on transcriptome data. BMC genomics. 24(1), 248 (2023).

